# Annotations capturing cell-type-specific TF binding explain a large fraction of disease heritability

**DOI:** 10.1101/474684

**Authors:** Bryce van de Geijn, Hilary Finucane, Steven Gazal, Farhad Hormozdiari, Tiffany Amariuta, Xuanyao Liu, Alexander Gusev, Po-Ru Loh, Yakir Reshef, Gleb Kichaev, Soumya Raychauduri, Alkes L. Price

## Abstract

It is widely known that regulatory variation plays a major role in complex disease and that cell-type-specific binding of transcription factors (TF) is critical to gene regulation, but genomic annotations from directly measured TF binding information are not currently available for most cell-type-TF pairs. Here, we construct cell-type-specific TF binding annotations by intersecting sequence-based TF binding predictions with cell-type-specific chromatin data; this strategy addresses both the limitation that identical sequences may be bound or unbound depending on surrounding chromatin context, and the limitation that sequence-based predictions are generally not cell-type-specific. We evaluated different combinations of sequence-based TF predictions and chromatin data by partitioning the heritability of 49 diseases and complex traits (average N=320K) using stratified LD score regression with the baseline-LD model (which is not cell-type-specific). We determined that 100bp windows around MotifMap sequenced-based TF binding predictions intersected with a union of six cell-type-specific chromatin marks (imputed using ChromImpute) performed best, with an 58% increase in heritability enrichment compared to the chromatin marks alone (11.6x vs 7.3x; P = 9 × 10^-14^ for difference) and a 12% increase in cell-type-specific signal conditional on annotations from the baseline-LD model (P = 8 × 10^-11^ for difference). Our results show that intersecting sequence-based TF predictions with cell-type-specific chromatin information can help refine genome-wide association signals.

## Introduction

Genome-wide association studies have revealed that non-coding genetic variation plays a central role in complex diseases and traits^1-3^; thus, understanding the syntax of non-coding genetic variation is of utmost importance. Transcription factors (TFs) are key elements of transcriptional regulation^4; 5^, and changes in their binding can ultimately affect human disease ^6-12^. Directly measuring TF binding is possible using ChIP-seq^13^; however, while TFs are numerous and their binding is cell-type-specific, ChIP-seq data has been generated for only a limited number of TFs and cell-types^14; 15^; A complete atlas of all TF binding sites would require tens of thousands of experiments, requiring immense resources. Many TFs bind specifically to unique motifs in the DNA sequence^16-18^ and their binding preferences can be inferred using sequence alone^19-27^. However, these sequence-based predictions often lack specificity as chromatin context has profound effects on TF binding. The vast majority of matches to a TF consensus sequence fall in regions of heterochromatin, which are inaccessible and therefore not actually bound^14^. It has been shown that incorporating open chromatin information from DNase-seq or ATAC-seq in addition to sequence can greatly improve prediction of TF binding^28; 29^. However, those methods use directly measured chromatin accessibility information and require learning footprints for each individual TF, making them difficult to apply to a diverse set of cell-types and factors. Moreover, they do not utilize functional information from histone modifications.

Here, we intersect various sequence-based TF annotations with cell-type specific chromatin annotations (including those imputed using ChromImpute^30^), creating cell-type-specific TF binding annotations for many tissues and cell-types. This strategy addresses both the limitation that identical sequences may be bound or unbound depending on surrounding chromatin context, and the limitation that sequence-based predictions are often not cell-type-specific. We use stratified LD score regression (S-LDSC) ^31^ with the baseline-LD model ^32^ to partition the heritability of 49 diseases and complex traits (average N=320K) in order to evaluate the contribution of these cell-type-specific TF binding annotations to disease.

## Results

### Cell-type-specific TF binding annotations recapitulate direct measurements of TF binding

To create more accurate annotations of cell-type-specific TF binding, we intersected sequence-based predictions with cell-type-specific chromatin annotations (Figure 1; see Supplemental Material and Methods). We constructed cell-type-specific annotations by taking the union of ChIP-seq peaks from five histone modifications that have previously been associated with active enhancers and promoters (H3K4me1, H3K4me2, H3K4me3, H3K9ac, and H3K27ac) as well as DNase1 Hypersensitive Sites (DHS) ^33^, available in 127 tissues and cell-types as part of the Roadmap Epigenomics project ^15^. Because experimental data is not available for every chromatin mark in every cell-type, we constructed two sets of annotations: one from all available directly measured peaks and one from imputed peaks computed for each chromatin mark and cell-type using ChromIMPUTE ^30^. We call these combined cell-type-specific chromatin annotations “Chromatin.measured” and “Chromatin.imputed”, respectively. We intersected the chromatin annotations with three sets of sequence-based TF binding predictions: MotifMap^19^, Kheradpour et al.^20^, and CisBP^34^. MotifMap uses sequence preferences from TRANSFAC ^16^ and JASPAR ^17; 18^ as well as conservation to predict binding. Kheradpour et al. trains many motif-finding methods on ENCODE TF ChIP-seq data and chooses those that perform best to apply genome-wide. CisBP is a large database of TF binding preferences from many sources. For each TF prediction set, we also tested annotations that include 20bp, 50bp, and 100bp windows. These windows may capture effects of sequence outside of the core motif^35^ and may capture cooperative binding sites for TFs that were not included in the datasets. We did not include sequence-based TF binding predictions produced by deep learning methods^22-26^ in our main analyses (see Discussion).

**Figure 1:**
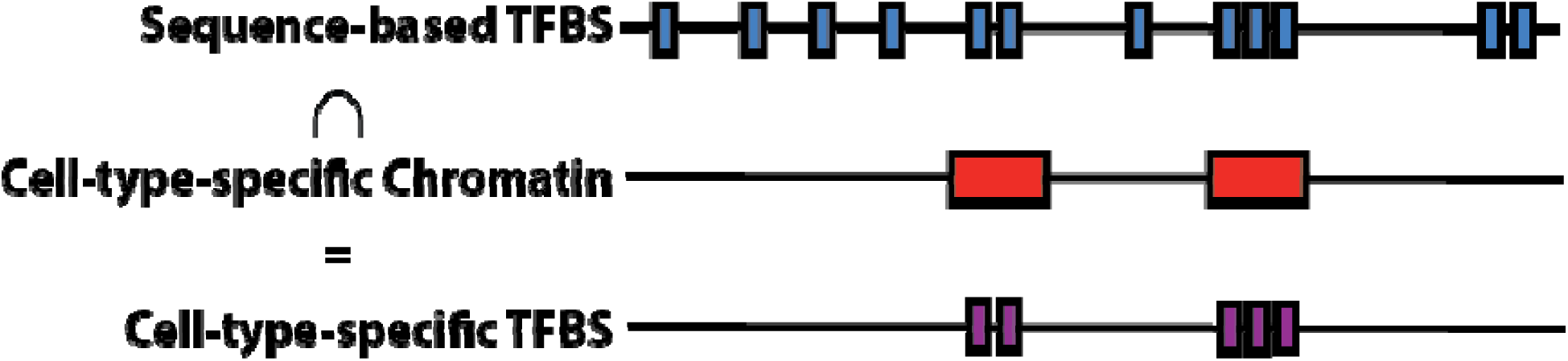
Strategy for constructing cell-type-specific TF binding annotations. We intersect sequence-based TF binding annotations such as MotifMap±100bp (blue bars; mean segment length 240bp) with cell-type-specific chromatin annotations (red bars; mean segment length 1200bp) to create cell-type-specific TF binding annotations such as Chromatinl1MotifMap100 (purple bars; mean segment length 220bp).

We assessed whether our new cell-type-specific TF binding annotations recapitulate direct measurements of TF binding. We compared ChIP-seq peaks from 91 experiments for 76 factors in lymphoblastoid cell lines (LCLs) from ENCODE ^14^ with the corresponding LCL-specific TF binding annotations and computed fold excess overlap (Figure 2 and Table S1; See Supplemental Material and Methods). As expected, there was only moderate excess overlap for the sequenced-based predictions: mean 1.69x (s.e. 0.02) across the three sequence-based predictions. However, the excess overlap was much larger when using either measured or imputed cell-type-specific chromatin annotations: 12.9x (s.e. 0.4) or 9.6x (s.e. 0.3) respectively. When the chromatin annotations were intersected with sequence-based predictions, the excess overlap increased, with the highest overlap in Chromatin.measured⍰CisBP: 17.6x (s.e. 0.7). Analysis of 5 other cell-types for which ChIP-seq TF binding data was available for at least 20 TFs produced similar conclusions (Table S1). This confirms that the new annotations are more accurately capturing cell-type-specific transcription factor binding. However, ChIP-seq peak may not provide a true gold-standard metric for capture of TF binding sites, as sequencing data peaks will also include regions surrounding the sites that are actually bound^36^. Moreover, ChIP-seq data is not available for many cell-type/TF pairs. We therefore turn to analysis of disease heritability to evaluate our annotations and investigate their potential applications.

**Figure 2:**
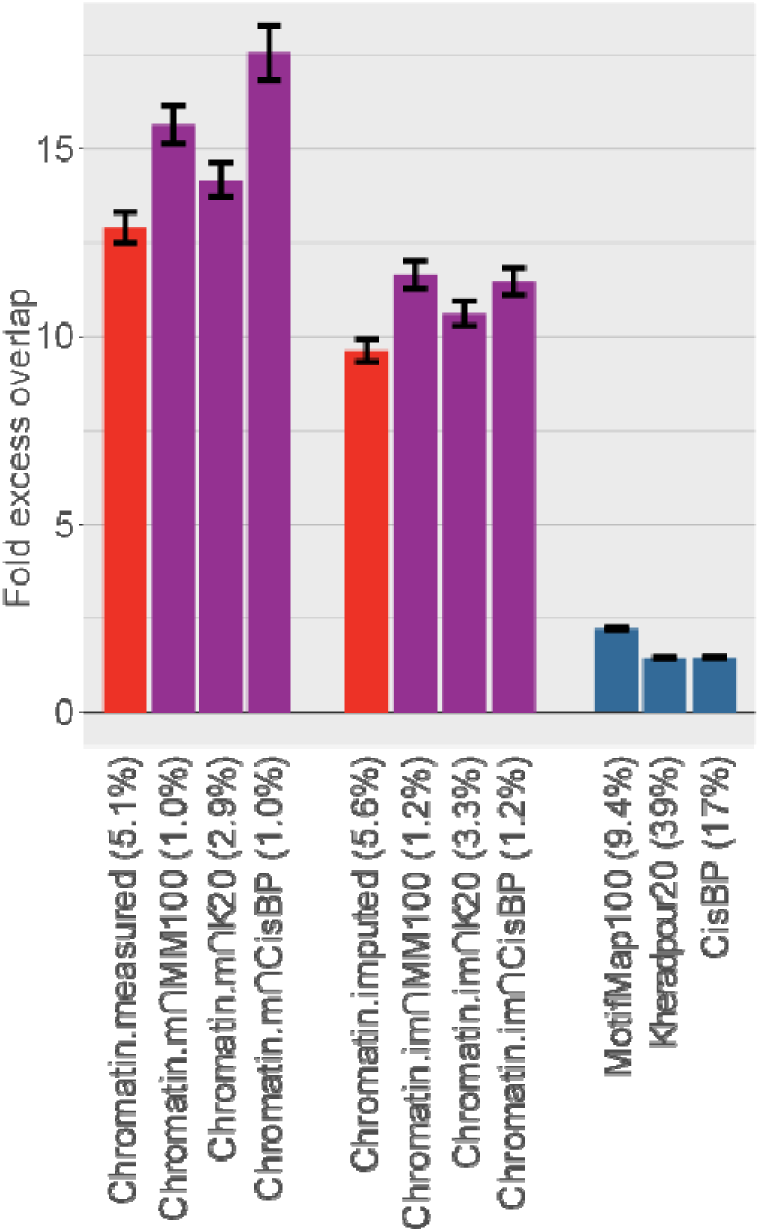
Comparison of excess overlap with TF ChIP-seq. We report the fold excess overlap with TF ChIP-seq peaks from ENCODE cell line GM12878 (lymphoblastoid cell line; data available from 91 experiments for 76 TFs) for sequence-based TF binding annotations (blue bars), cell-type-specific chromatin annotations (red bars), and cell-type specific TF binding annotations (purple bars). Error bars denote one standard error. The percentage under each bar indicates the proportion of SNPs in each annotation. Numerical results, including results for 5 other tissues for which ChIP-seq TF binding data was available for at least 20 TFs, are reported in Table S1.

### Cell-type-specific TF binding annotations are enriched for disease heritability

We assessed whether our new cell-type-specific TF binding annotations are enriched for disease heritability. We used two metrics to quantify the contribution of an annotation to disease heritability: enrichment and standardized effect size (τ*) (see Supplemental Material and Methods). Enrichment is defined as the proportion of heritability explained by SNPs in an annotation divided by the proportion of SNPs in the annotation^31^. τ* is defined as the proportionate change in per-SNP heritability associated with an increase in the value of the annotation by one standard deviation, conditional on other annotations included in the model^32^. Unlike enrichment, τ* quantifies effects that are unique to the focal annotation.

We analyzed ^49^ diseases and complex traits for which summary association statistics are publicly available (Table S2; average N=320k), and analyzed 127 Roadmap tissues and cell-types ^15^. For each (trait,cell-type) pair, we ran stratified LD score regression (S-LDSC) ^31^ using the baseline-LD model v2.0 (^32^; see Web Resources) and the corresponding cell-type-specific chromatin annotation. For each trait, we chose the best cell-type based on statistical significance of τ* for the cell-type-specific chromatin annotation, consistent with previous work ^31^ (Table S2). We used this cell type for all cell-type-specific annotations for that trait; this is a conservative choice when comparing our new cell-type-specific TF binding annotations to cell-type-specific chromatin annotations.

We sought to identify the most disease-informative way to combine sequence-based TF binding predictions and cell-type-specific chromatin annotations. For each combination of 24 cell-type-specific TF binding annotations (3 sequence-based TF predictions × 4 window sizes [0bp,20bp,50bp,100bp] × 2 chromatin types [measured,imputed]), we ran S-LDSC conditional on the baseline-LD model and the cell-type-specific chromatin annotation. We meta-analyzed results across the 49 traits and calculated three metrics for each annotation: heritability enrichment, τ*, and combined τ*; combined τ* is a generalization of τ* that quantifies the combined information in the cell-type-specific chromatin and cell-type-specific TF binding annotations, conditional on the baseline-LD model (see Supplemental Material and Methods).

Results of the meta-analysis across 49 traits are reported in Figure 3 (6 cell-type-specific TF binding annotation s; 3 sequence-based TF predictions × 2 chromatin types, with best window size for each) and Table S3. We first note that imputed chromatin consistently attained slightly smaller heritability enrichment but slightly higher τ* than measured chromatin; since the imputed chromatin data is complete for all 127 Roadmap cell-types, we focused on imputed chromatin for subsequent analyses. The Chromatin⍰MotifMap100 annotations performed best, with a 59% higher heritability enrichment than Chromatin (11.6x vs 7.3x, p=9 × 10^-14^ for difference) and a 12% higher combined τ* (1.87 vs 1.67 τ*; p=8 × 10^-11^ for difference); these improvements are statistically significant after correcting for 24 hypotheses tested. The τ* values, reflecting information unique to these annotations, were very large relative to analogous values (τ* up to 0.52) that we recently estimated for non-cell-type-specific LD-related annotations ^32^ and molecular QTL annotations ^37^; as such, the Δτ* of 0.20 is a substantial improvement. Chromatin⍰Kheradpour^20^ and Chromatin⍰CisBP attained slightly worse results. Unsurprisingly, the sequence-based TF binding annotations alone attained relatively low enrichment and τ*.

**Figure 3:**
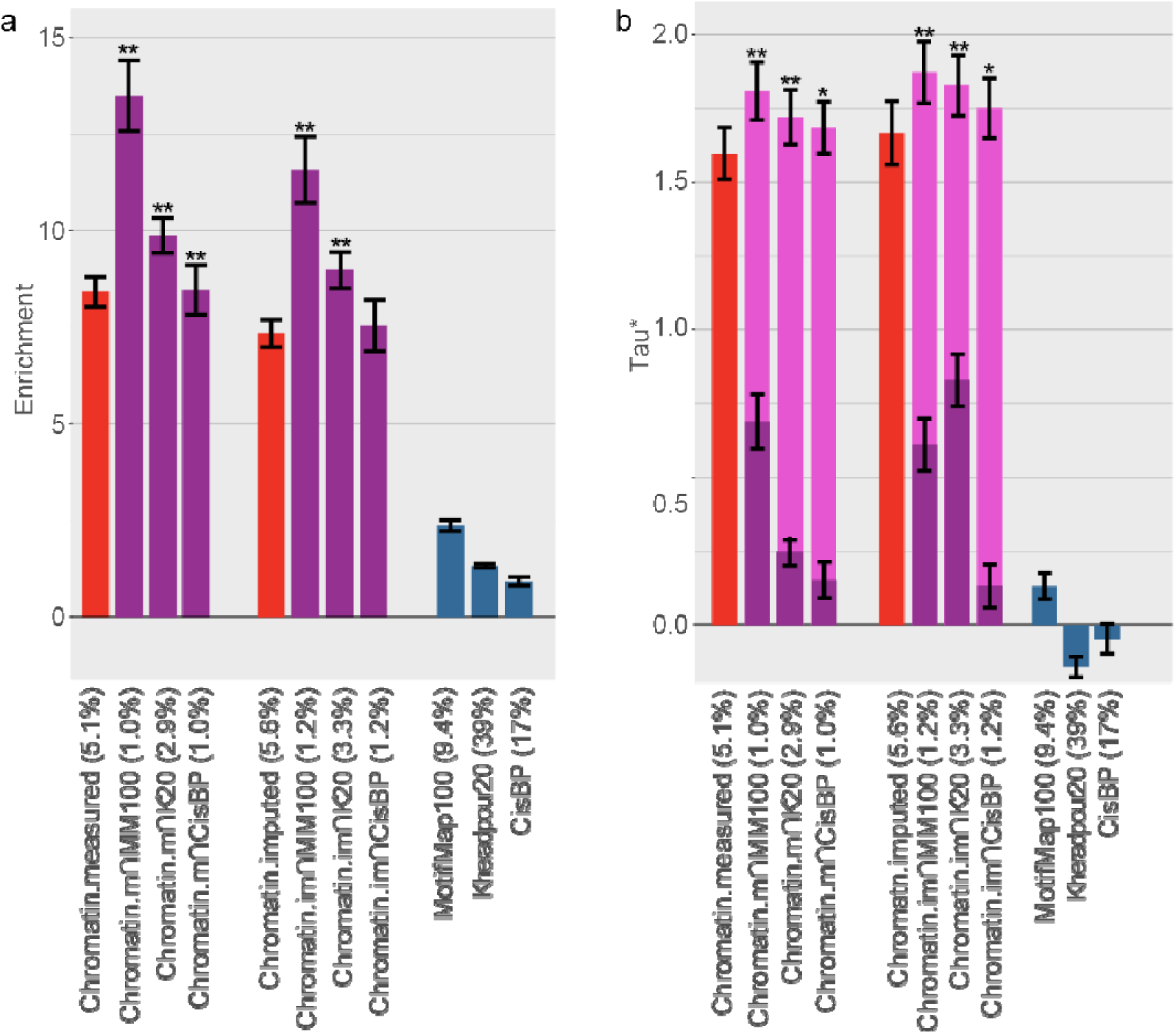
Comparison of heritability enrichment and τ* across 49 diseases and complex traits. We report (a) heritability enrichment and (b) τ* for sequence-based TF binding annotations (blue bars), tissue-specific chromatin annotations (red bars), and tissue-specific TF binding annotations (purple bars, including dark purple bars for τ* in a joint model and light purple bars for combined τ*). Error bars denote one standard error. (*) p<0.05 for (a) enrichment vs corresponding chromatin annotation and (b) combined τ* vs. τ* of corresponding chromatin annotation. (**) p-value < 1e-5. The percentage under each bar indicates the proportion of SNPs in each annotation. Numerical results are reported in Table S3 and S4.

Results of targeted meta-analyses across 6 autoimmune, 5 blood, and 11 brain-related traits (See Supplemental Material and methods) are reported in Figure 4 and Table S4. For the 6 autoimmune traits, Chromatin⍰MotifMap100 attained a much higher heritability enrichment than Chromatin (23.6x vs. 11.1x; p=0.001 for difference) and a substantially higher combined τ* (3.11 vs. 2.61; p=0.004 for difference). Chromatin⍰MotifMap100 also outperformed TF-binding annotations from ENCODE ChIP-seq (heritability enrichment=23.6x vs. 5.32x; τ*=3.11 vs. 1.75). Results were similar for the 5 blood traits, though enrichments were slightly smaller and differences less significant. On the other hand, the 11 brain-related traits attained substantially smaller enrichments, consistent with previous work ^31; 32^. However, Chromatin⍰MotifMap100 still attained substantial improvements in heritability enrichment (7.04 vs. 4.98; p=0.002 for difference) and τ* (1.17 vs 1.07; p=0.003 for difference). We also considered an annotation constructed from the union of all ENCODE ChIP-seq TF binding experiments. Notably, this annotation underperformed Chromatin⍰MotifMap100 for all trait classes, and performed particularly poorly for the brain-related traits (heritability enrichment=1.31, τ*=0.07). This is likely because very few of the ENCODE ChIP-seq experiments were conducted in brain tissues, highlighting the importance of methods to create cell-type-specific TF annotations when ChIP-seq data is unavailable.

**Figure 4:**
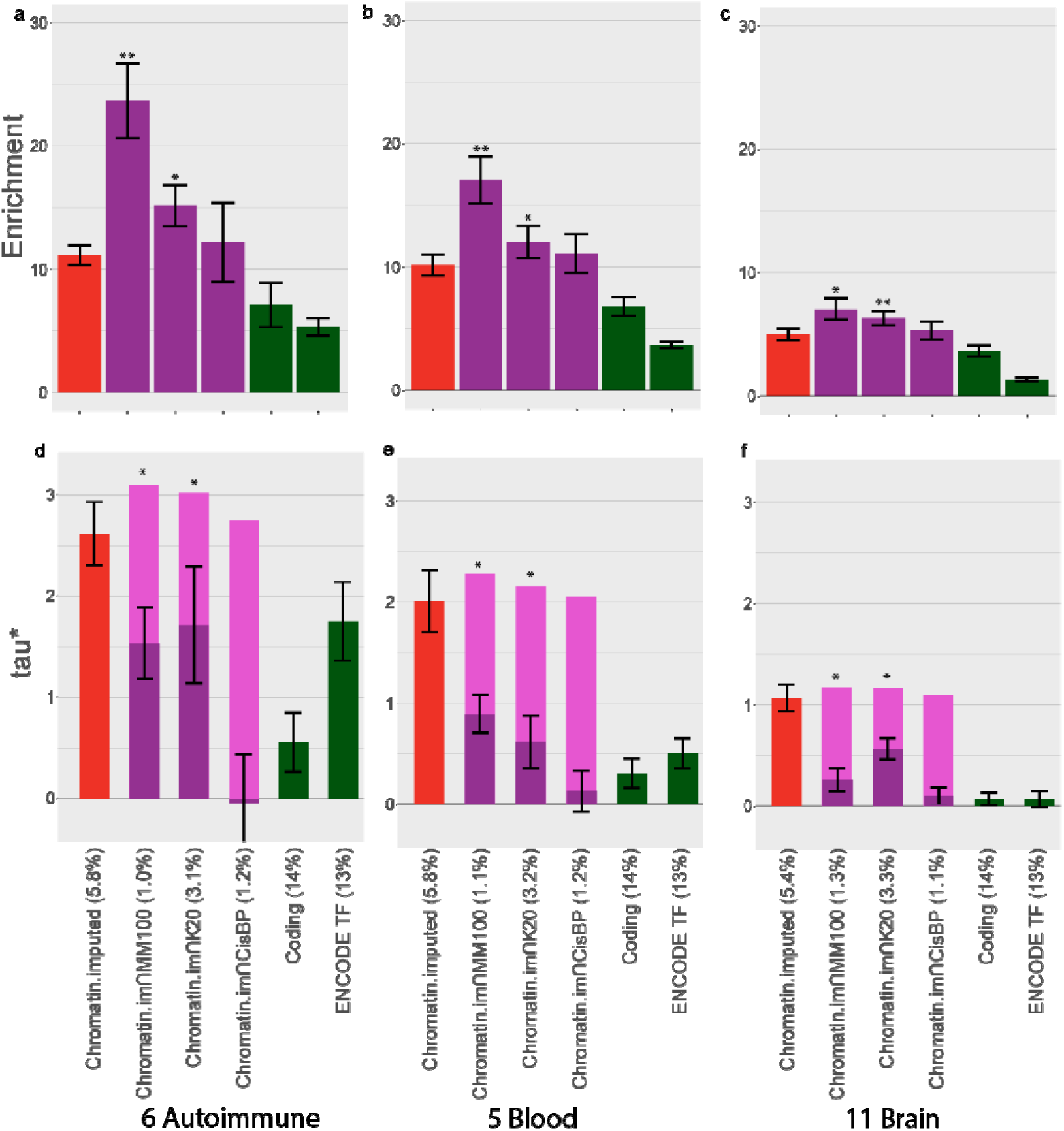
Comparison of heritability enrichment and τ* for autoimmune, blood and brain-related traits. We report (a-c) heritability enrichment for each trait class and (d-f) τ* for each trait class for cell-type-specific chromatin annotations (red bars), cell-type-specific TF binding annotations (purple bars, including dark purple bars for τ* in a joint model and light purple bars for combined τ*). We include coding regions (green bars) and an annotation constructed from the union of all ENCODE ChIP-seq TF binding experiments (green bars) for comparison purposes. Error bars denote one standard error. (*) p<0.05 for (a) enrichment vs corresponding chromatin annotation and (b) combined τ* vs. τ* of corresponding chromatin annotation. (**) p-value < 1e-5. The percentage under each bar indicates the proportion of SNPs in each annotation. Numerical results are reported in Table S5 and S6.

Finally, we compared the heritability enrichments of Chromatin and Chromatin⍰MotifMap100 for each individual trait (Figure 5 and Table S5). We determined that 43 of 44 traits with significant enrichment for at least one of these two annotations had higher heritability enrichment for Chromatin⍰MotifMap100. However, some traits show only modest improvements in heritability enrichment, perhaps because binding preferences for the relevant TFs are not well-captured by sequence-based predictions; alternatively, it is possible that TF binding sites play smaller roles for these traits.

**Figure 5:**
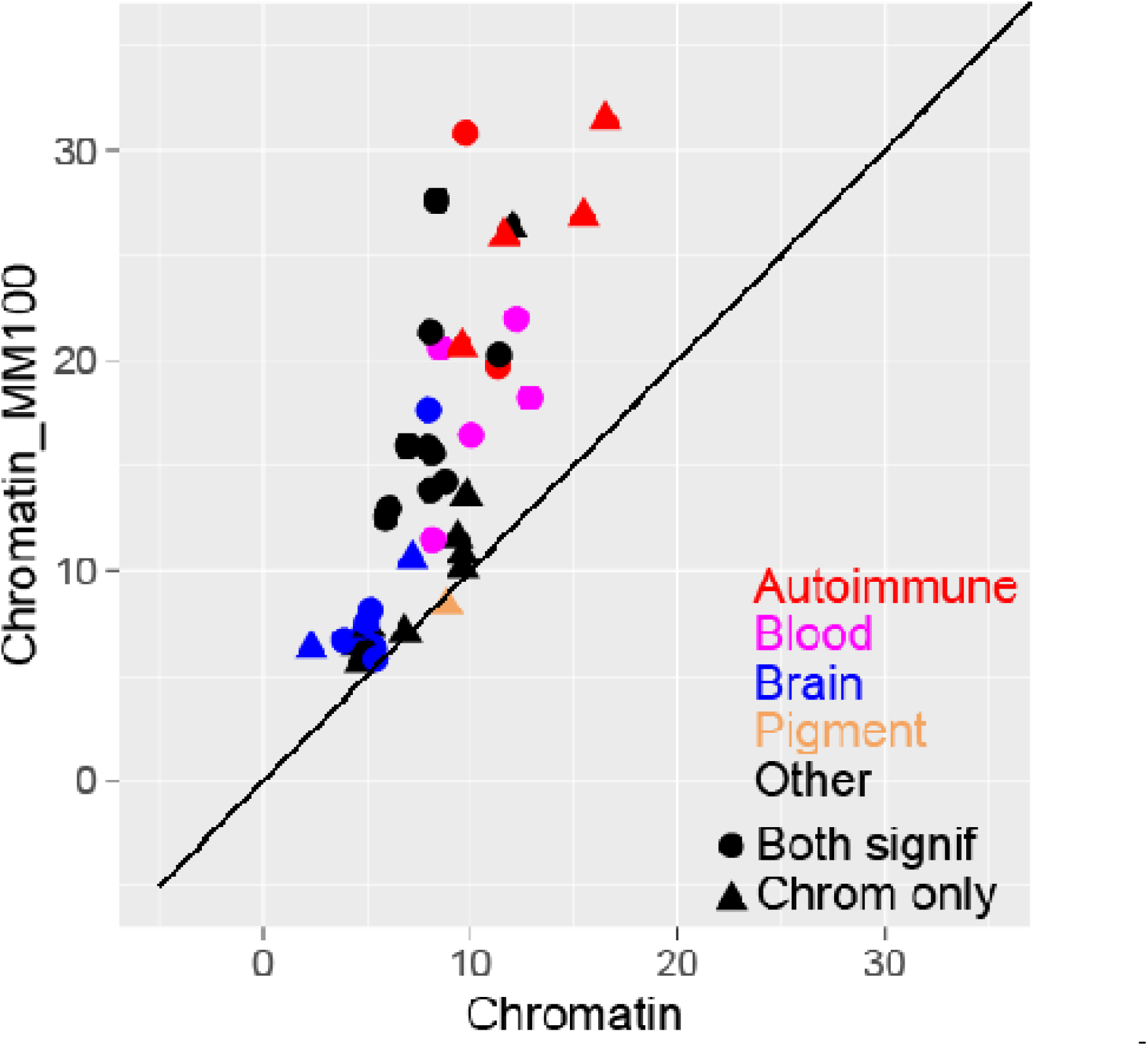
Comparison of heritability enrichment for each trait. We report the heritability enrichment of cell-type-specific chromatin annotations (x-axis) and cell-type-specific TF binding annotations (y-axis). Results are displayed for 44 traits that have significant enrichment for at least one of these two annotations, assessed using p=0.05/127, (correcting for 127 cell-types analyzed). Numerical results are reported in Table S7.

### Choice of baseline vs. baseline-LD model in cell-type-specific analyses

Our main analyses (Figures 3-5) used the baseline-LD model (v2.0), which includes 6 LD-related annotations ^32^; using a more complete model is appropriate when the goal is to estimate heritability enrichment while minimizing bias due to model misspecification ^31; 32^. On the other hand, our recent work ^31; 38^ identified critical cell types for disease by computing the statistical significance of τ* conditioned on the baseline model, which does not include the LD-related annotations. The LD-related annotations reflect the action of negative selection ^32^; some of the LD-related annotations are correlated with cell-type-specific annotations—particularly brain annotations, which show stronger signals of negative selection ^39^. Thus, we hypothesized that cell-type-specific signals might be stronger when conditioning on the baseline model instead of the baseline-LD model. To assess this, we compared the statistical significance of the combined τ* for (Chromatin + Chromatin⍰MotifMap100) using the baseline (v1.1) vs. baseline-LD (v2.0) models across 49 traits; in each case, we chose the most significant of the 127 Roadmap cell types. We determined that the baseline model generally produces more significant combined τ* values than the baseline-LD model, particularly for brain traits and cell types (Figure 6 and Table S6). Thus, we recommend that the baseline model should be used when the goal is to identify critical cell types; however, the baseline-LD model should still be used when the goal is to obtain unbiased estimates of heritability enrichment.

**Figure 6:**
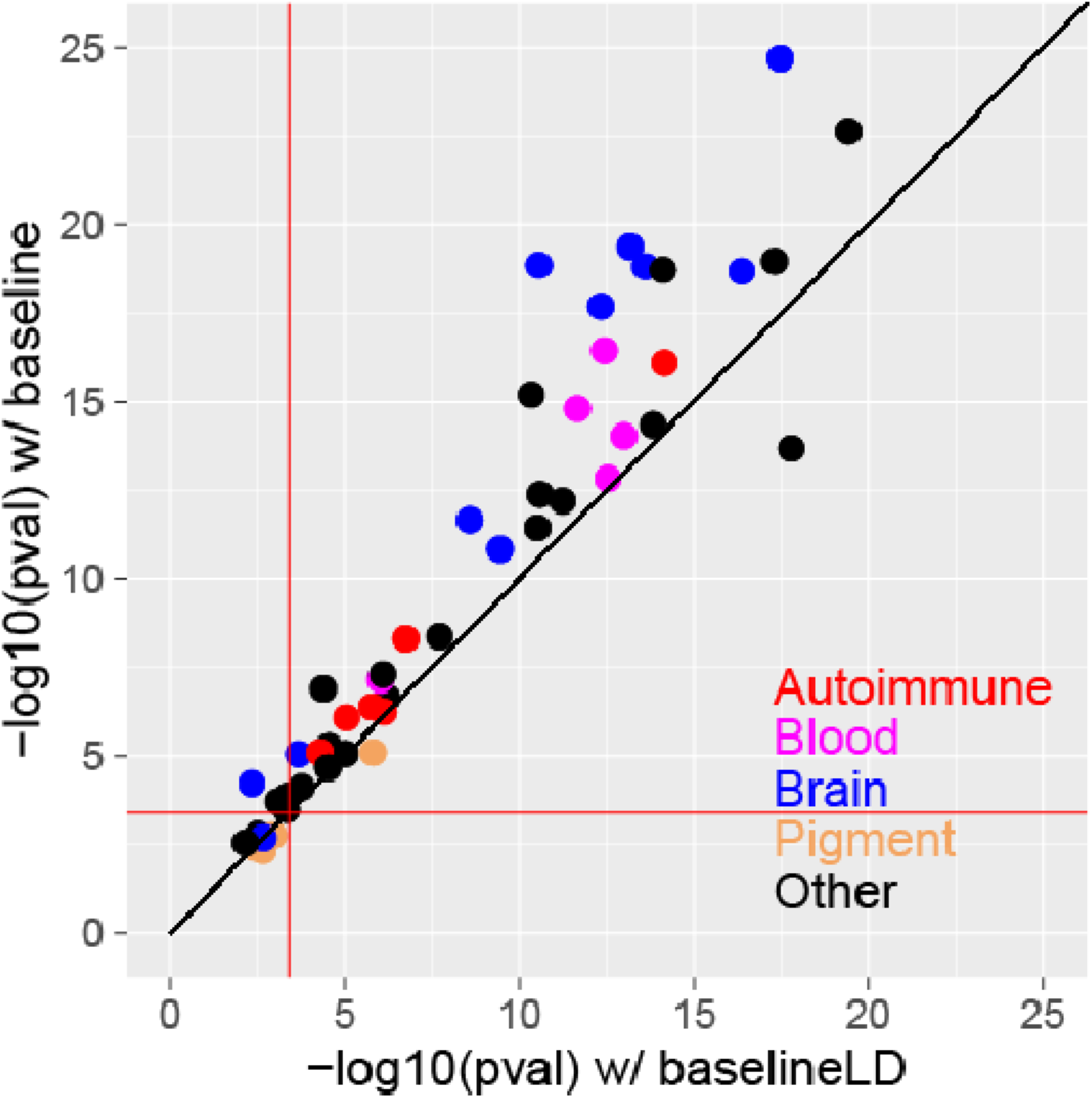
Comparison of combined cell-type-specific annotations (Chromatin + Chromatin[lMotifMap100) conditioned on the baseline vs. baseline-LD models. We report the statistical significance (l1log10P-value of combined τ*) of the combined cell-type-specific annotations (Chromatin + Chromatinl1MotifMap100) for the baseline (y-axis) vs. baseline-LD (x-axis) models, for each of 49 traits. In each case, we report results for the most significant tissue/cell type. The red lines indicate the p=0.05/127 significance threshold, correcting for testing of 127 cell-types. Numerical results are reported in Table S8.

## Discussion

We explored a new strategy for constructing cell-type-specific TF binding annotations by intersecting sequence-based TF predictions with cell-type-specific chromatin annotations. We determined that the resulting cell-type-specific TF binding annotations significantly outperformed cell-type-specific chromatin annotations across 49 diseases and complex traits, with highly significant improvements in both heritability enrichment and τ*; this strategy increased heritability enrichment for 43 of 44 traits with significant conditional signal for cell-type-specific chromatin, and greatly outperformed non-cell-type-specific sequence-based TF binding annotations. These findings are consistent with the higher overlap of our cell-type-specific TF binding annotations with ENCODE TF ChIP-seq peaks. We also determined that annotations constructed using imputed chromatin^30^ attained slightly higher τ* than annotations constructed using measured chromatin; we recommend the use of imputed chromatin annotations, since they are complete for all 127 Roadmap cell-types.

Our results confirm that TF binding is important for diseases and complex traits, and provide a quantification of their contribution to heritability. In particular, a large proportion of the heritability explained by active chromatin regions comes from predicted TF binding sites, particularly for autoimmune diseases. This proportion will only increase as our TF predictions improve. Therefore, we recommend that cell-type-specific TF binding annotations should be incorporated into efforts to interpret GWAS signals using functional fine-mapping^3; 21; 40; 41^, as well as efforts to use functional information to increase association power^42-44^ and improve polygenic risk prediction ^45-47^.

We note three limitations of our work. First, our cell-type-specific TF binding annotations attain higher heritability enrichment than cell-type-specific chromatin annotations, but explain less heritability in total due to their smaller size. We evaluated this tradeoff using the τ* metric ^32^, which demonstrated that our cell-type-specific TF binding annotations attain a highly significant increase in cell-type-specific signal conditional on the baseline-LD model, compared to cell-type-specific chromatin annotations alone. Second, we did not include sequence-based TF binding predictions produced by deep learning methods in our main analyses ^23-26^. The reason for this is that combining annotations across TFs adds an additional layer of complexity, as TF binding predictions for different TFs are not on the same scale; in particular, TF consensus sequences vary in size and the number of sites bound by a TF varies greatly. We investigated several strategies for combining TF binding predictions produced by DeepBind^24^ across TFs, but we were unable to devise a strategy that attained performance close to the strategies that we report here (Figure 3). Third, the sequence-based predictions that we incorporate are limited to TFs that have sufficient data available to learn the underlying consensus sequence. It is possibly that TFs active in some cell-types (e.g. skin) are underrepresented, potentially explaining why some traits (e.g. pigmentation traits) perform less well in our analyses. Fourth, inferences about components of heritability can potentially be biased by failure to account for LD-dependent architectures^32; 48-50^. All of our main analyses used the baseline-LD model, which includes 6 LD-related annotations^32^. The baseline-LD model is supported by formal model comparisons using likelihood and polygenic prediction methods, as well as analyses using a combined model incorporating alternative approaches^51^; however, there can be no guarantee that the baseline-LD model perfectly captures LD-dependent architectures. Despite these limitations, our tissue-specific TF binding annotations significantly improve our understanding of disease and complex trait heritability. All annotations have been made publicly available (see Web Resources).

## Acknowledgements

We are grateful to Manolis Kellis, Yue Li, and Babak Alipanahi for helpful discussions. This research was funded by NIH grants U01 HG009379, R01 MH101244, R01 MH109978, R01 MH107649 and F32 HG009615, and by a McLennan Family Fund award. This research was conducted using the UK Biobank Resource under Application 16549.

## Supplemental Material and Methods

### Constructing sequence-based TF binding annotations

MotifMap: Predicted TF binding sites for build hg19 were downloaded from the MotifMap website (http://motifmap.igb.uci.edu)

Kheradpour et al.: Predicted TF binding sites for build hg19 were downloaded from http://compbio.mit.edu/encode-motifs/matches.txt.gz

CisBP: Position weight matrixes for all human TFs were downloaded from the CisBP website (http://cisbp.ccbr.utoronto.ca/). Genome-wide matches were created using MEME FIMO software (http://meme-suite.org/doc/fimo.html), which provides p-values for the match of a given sequence to a motif above the background genomic sequence. Matches with p-value < 1e-5 for each TF were kept.

DeepBind: We downloaded DeepBind and the human TF models from the DeepBind website (http://tools.genes.toronto.edu/deepbind). We then constructed fasta files spanning the entire genome with overlapping 101 base pair lines of sequence as input for a genome-wide DeepBind scan. We ran DeepBind genome-wide for each TF as well as on a gold-standard set of sequence from ChIP-seq data (also downloaded from the DeepBind site). We then assigned each 101 base pair line a z-score for binding based on (1) the mean and standard deviation of the gold-standard sequences or (2) the mean and standard deviation of the genome-wide scores. We constructed binding annotations using various thresholds for both, but no combination yielded positive results.

### Assessing overlap with direct measurements of ChIP-seq

We downloaded TF ChIP-seq data from ENCODE (http://hgdownload.cse.ucsc.edu/goldenpath/hg19/encodeDCC/wgEncodeAwgTfbsUniform/). For each cell-type with at least 30 experiments, we created a bed file with the union of ChIP-seq peaks from all TFs assayed. We then calculated the excess overlap between an annotation (A) and the ChIP-seq peaks (B) as

(1)

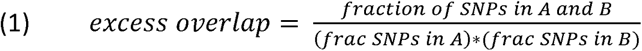

We calculated standard errors using a block-jackknife, dividing the genome into 200 blocks of equal genomic size.

### Choosing best cell-type for each disease

We applied S-LDSC conditional on the baseline-LD model (v2.0) with “Chromatin.imputed” annotations for each pair of 127 Roadmap cell-types and 49 traits. We then chose the most disease-relevant cell-type based on significance of τ*.

### Calculating combined τ*

In order to calculate combined τ*, we applied S-LDSC conditional on the baseline-LD model and including both Chromatin and one Chromatinl1TFBS annotation at a time. We then calculated

(2)

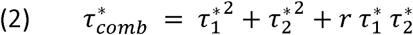

where 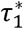 and 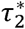 are the τ* for Chromatin and Chromatinl1TFBS respectively and r is the correlation between the Chromatin and Chromatinl1TFBS annotations. We calculated standard errors for 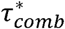 using a block-jackknife with 200 blocks. We also calculated p-values for the difference between 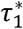 and 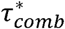 by jackknifing on the value (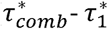). With this metric, we measure the combined information being captured by a set of cell-type-specific annotations.

## Web Resources

CisBP, http://cisbp.ccbr.utoronto.ca/

DeebBind, http://tools.genes.toronto.edu/deepbind

ENCODE, http://hgdownload.cse.ucsc.edu/goldenpath/hg19/encodeDCC

Kheradpour et al., http://compbio.mit.edu/encode-motifs

LDSC software, https://github.com/bulik/ldsc/wiki

LDSC annotations, https://data.broadinstitute.org/alkesgroup/LDSCORE/ MEME, http://meme-suite.org/doc/fimo.html

**See attached excel file**

**Table S1: Comparison of excess overlap with TF ChIP-seq.** We report the fold excess overlap with TF ChIP-seq peaks from 6 cell-types from ENCODE. For each cell-type, we match with the corresponding Roadmap cell-type and test sequence-based TF binding annotations, cell-type-specific chromatin annotations, and cell-type specific TF binding annotations.

**See attached excel file**

**Table S2: Choice of best cell-type for each trait.** We report the fold excess overlap with TF ChIP-seq peaks from 6 tissues and cell lines from ENCODE. We test sequence-based TF binding annotations, cell-type-specific chromatin annotations, and cell-type specific TF binding annotations.

**See attached excel file**

**Table S3: Comparison of heritability enrichment and τ* across 49 diseases and complex traits.** (A) We report heritability enrichment as well as standard errors for each annotation. Enrichments are meta-analyzed across 49 traits. (B) We report τ* and standard error for each cell-type-specific chromatin annotation fit independently with the baseline-LD model and meta-analyzed across 49 traits. We then report τ* and standard error for each cell-type-specific TF binding annotation fit with the baseline-LD model and the corresponding chromatin annotations. We also report combined τ* for each cell-type specific TF binding annotation as well as the difference between combined τ* and the τ* of the chromatin annotation alone.

**See attached excel file**

**Table S4: Comparison of heritability enrichment and τ* for autoimmune, blood and brain-related traits.** (A) We report heritability enrichment as well as standard errors for each annotation. Enrichments are meta-analyzed across 6 autoimmune, 5 blood, and 11 brain-related traits respectively. (B) We report τ* and standard error for each cell-type-specific chromatin annotation fit independently with the baseline-LD model and meta-analyzed for 6 autoimmune, 5 blood, and 11 brain-related traits respectively. We then report τ* and standard error for each cell-type-specific TF binding annotation fit with the baseline-LD model and the corresponding chromatin annotations. We also report combined τ* for each cell-type specific TF binding annotation as well as the difference between combined τ* and the τ* of the chromatin annotation alone.

**See attached excel file**

**Table S5: Comparison of heritability enrichment for each trait.** We report the heritability enrichment of cell-type-specific chromatin annotations and cell-type-specific TF binding annotations.

**See attached excel file**

**Table S6. Comparison of combined cell-type-specific annotations (Chromatin + ChromatinllMotifMap100) conditioned on the baseline vs.** baseline-LD models. We report the statistical significance of the combined cell-type-specific annotations (Chromatin + Chromatinl1MotifMap100) for the baseline vs. baseline-LD models, for each of 49 traits.

